# Adaptive resistance to FLT3 inhibitors is potentiated by ROS-driven DNA repair signalling

**DOI:** 10.1101/2024.07.26.605229

**Authors:** Dilana E. Staudt, Zacary P. Germon, Abdul Mannan, Tabitha McLachlan, Heather C. Murray, Ryan J. Duchatel, Bryce C. Thomas, Tyrone Beitaki, Holly P. McEwen, Mika L. Persson, Leah Calvert, Izac J. Findlay, Evangeline R. Jackson, Nathan D. Smith, David A. Skerrett-Byrne, David Mossman, Brett Nixon, Geoffry De Iullis, Alicia M. Douglas, Anoop K. Enjeti, Jonathan R. Sillar, Janis Chamberlain, Frank Alvaro, Andrew H. Wei, Patrick Connerty, Nicole Verrills, Matthew D. Dun

## Abstract

Alterations in the FMS-like tyrosine kinase 3 (FLT3) gene are the most frequent driver mutations in acute myeloid leukaemia (AML), linked to a high risk of relapse in patients with internal tandem duplications (FLT3-ITD). Tyrosine kinase inhibitors (TKIs) targeting the FLT3 protein are approved for clinical use, yet resistance often emerges. This resistance is mainly seen following the acquisition of additional point mutations in the tyrosine kinase domain (TKD), resulting in a double mutant FLT3-ITD/TKD, which sustains cell signalling and survival despite the presence of FLT3 inhibitors. Here, we developed a FLT3-mutant AML model with adaptive resistance to type II TKIs, sorafenib, and quizartinib by *in vitro* drug selection. Through global multiomic profiling, we identified upregulation of proteins involved in reactive oxygen species (ROS) production, particularly NADPH-oxidases, driving cellular ’ROS-addiction’, with resistant cells relying on ROS for survival, and genome fidelity preserved by ATM-driven DNA repair. Transcriptomic analysis of adult and paediatric AML (pAML) patients identified high ATM expression as a biomarker for shorter median overall survival in both the de novo and relapsed settings. Inhibition of ATM with clinically relevant therapy WSD-0628 effectively killed TKI- and chemotherapy-resistant AML cells *in vitro* and significantly extended the survival of mice with sorafenib- and quizartinib-resistant FLT3-ITD AML *in vivo*. We propose a new treatment strategy to improve survival of patients who develop resistance to sorafenib and quizartinib, as well as relapsed and refractory pAML, exploiting resistance mechanisms to precision therapies and cell-intrinsic features of high-risk cases, highlighting a clinically relevant salvage strategy.

## INTRODUCTION

Constitutive activation of the FMS-like Tyrosine Kinase 3 receptor (FLT3; CD135) following the acquisition of genetic mutations is seen in 20-30% of acute myeloid leukaemia (AML) cases (1), leading to high relapse risk and poor survival (2, 3). FLT3, a type III receptor tyrosine kinase (RTK-III), regulates haematopoietic cell differentiation, proliferation, and survival (4). Binding of FLT3 ligand (FLT3L) activates ERK/MAPK (5), JAK/STAT (6), and PI3K/AKT signalling (7). The most common FLT3 mutations in AML are internal tandem duplications (FLT3-ITD; 20-25%) (8), followed by point mutations in the second tyrosine kinase domain (FLT3-TKD; 5-10%) (8, 9) both causing constitutive FLT3 activation and oncogenic signalling (9).

The high prevalence of FLT3 mutations in AML has led to the development and clinical assessment of first- and second-generation tyrosine kinase inhibitors (TKIs) (1, 10, 11). However, single-agent TKI therapies often fail long-term due to unsustained anti-leukaemic responses and secondary resistance (10–12). Midostaurin (PKC412) was the first TKI approved for newly diagnosed FLT3-mutant AML with standard cytarabine and anthracycline therapy in 2017 (13). Quizartinib (AC220), a second-generation FLT3 inhibitor, was approved in 2023 for use with standard therapy and as maintenance monotherapy (14). Gilteritinib is approved for relapsed/refractory AML (R/R-AML) with FLT3 mutation (15). Several other TKIs are in clinical assessment for AML (Supplementary Table S1).

Sorafenib, initially approved for solid tumours, shows promise in FLT3-ITD AML as a maintenance therapy post-allo-HCT, improving relapse-free survival and as salvage therapy for post-allo-HCT relapses (16). It is extensively studied for FLT3-mutant paediatric AML (pAML) (17), offering multi-kinase inhibition, including FLT3, to combat resistance and improve outcomes.

Despite initial improvements, resistance to FLT3-targeted therapies persists (10–12), caused by FLT3L overexpression (18), clonal selection (19), bone marrow stromal protection (20), off-target mutations, FLT3-independent/ downstream signalling pathway activation (21), or through additional mutations in FLT3 (FLT3-ITD/TKD) affecting drug binding sites (1, 10, 22, 23). Mutations at residues D835 (10, 12, 24), Y842 (22, 23), F691 (23), N676 (22), and A627 (23) confer resistance to type I and/or II TKIs.

Understanding oncogenic signalling pathways active in high-risk and resistance settings aids precision medicine. This study integrates next-generation sequencing (NGS), phosphoproteomics, and patient transcriptomics to identify upregulation of the oxidative stress-driven ATM DNA repair signalling pathway in therapy-resistant AML, effectively targeted by the novel ATM inhibitor, WSD-0628. These findings support early-phase trials for R/R AML or high-risk pAML.

## MATERIALS AND METHODS

Detailed materials and methods can be found in the Supplemental Information.

### Study approval

The use of patient-derived human AML cell lines was approved by the Human Ethics Research Committee, University of Newcastle (H-2018-0241). All *in vivo* studies were approved by the University of Newcastle Animal Care and Ethics Committee (A-2017-733, A-2023-308).

### In vitro development of adaptive resistance

TKI and standard-of-care resistant MV4-11 cells were developed through serial passaging of cells in increasing doses of sorafenib (Selleckchem, Houston, TX, USA) from 2.5 nM-1280 nM over 20 weeks or cytarabine (Selleckchem) from 10 µM-500 µM followed by daunorubicin (Sigma-Aldrich, Burlington, MA, USA) from 2 nM-12 nM. Cells were DNA sequenced by next generation sequencing (Supplementary Table S2) using the Myeloid Solution panel, as described in Supplemental Information.

### Structural modelling of tyrosine kinase inhibitors binding to FLT3

Several crystal structures detailing the intracellular domains of FLT3, including the inactive conformation of the TKDs were used as templates to create a multiple sequence alignment homology model of the inactive kinase as previously described (25).

### pHASED phosphoproteomics

Quantitative phosphoproteomics of MV4-11 parental and TKI resistant cells was performed as previously described (26). The mass spectrometry proteomics data is deposited to the ProteomeXchange Consortium via the PRIDE partner repository (27) with the dataset identifier PXD053329 and 10.6019/PXD053329. Reviewer access via the PRIDE website - **Username:** **Password:** lTeh0PPlMcFj

### Analysis of publicly available patient survival and expression data

RNA-seq and clinical data from the Therapeutically Applicable Research to Generate Effective Treatments (TARGET) and Beat AML initiatives was minded and analysed using GraphPad Prism Software (version 10.0.2, GraphPad, Boston, MA, USA) (28).

### Detection of reactive oxygen species

Dihydroethidium (DHE) (Life Technologies, Australia) was used to detect intracellular cytoplasmic superoxide as previously described (29).

### ATM knockdown

MV4-11 sensitive and resistant cell lines were transfected using RNAiMAX lipofectamine reagent (Invitrogen, Carlsbad, CA) as per manufacturer’s instructions with ATM–specific small interfering RNA (siRNA) or a scrambled siRNA control.

*Terminal Deoxynucleotidyl Transferase dUTP Nick End Labelling (TUNEL) DNA Fragmentation Assay and Oxidative DNA Damage Assessment by 8-hydroxy-2′-deoxyguanosine (8-OHdG)* MV4-11 FLT3-ITD sensitive and TKI resistant cell lines were treated with either 250 nM WSD-0628 for 24 h, 1 mM H2O2 for 30 min or ATM-siRNA or scramble control and assessed for formation of TUNEL or oxidative DNA damage through 8-OHdG ICC fluorescence (30).

### AML xenograft mouse modelling

NOD-*Rag1^null^ IL2rg^null^* (NRG) mice were engrafted with FLT3-ITD TKI sensitive or resistant, MV4-11-luc cells (1e^6^ cells in PBS) by tail vein injection. Bioluminescence imaging (BLI) was used to detect leukemic cell engraftment, and randomized for treatment once BLI reached a mean radiance of 1 × 10^6^ p/s. Mice were treated with either vehicle control, sorafenib (10 mg/kg/day MV4-11 resistant; 2.5 mg/kg/day MV4-11 sensitive), quizartinib (2 mg/kg/day) or WSD-0628 (5 mg/kg/day) as monotherapies, or in combination, for four weeks. Mice were euthanized at ethical endpoint, including weight loss exceeding 20% or body condition scores indicating ethical endpoint.

## RESULTS

### Establishment of a human cell line model of adaptive TKI resistance

Dual FLT3-ITD and D835V/Y mutations are a known mechanism of resistance to type II TKIs. Previously, we generated isogenic FLT3-ITD AML cell line models with these mutations (24). Quantitative proteomics identified the activation of the ATM pathway, promoting cell survival despite high dose sorafenib (26). Here, we have expanded these studies by developing a human FLT3-ITD AML cell line (MV4-11) model of adaptive resistance to type II TKIs by treating cells with increasing sorafenib doses (0 nM - 1280 nM) (Supplementary Figure S1). Next generation sequencing (NGS) was performed on the sorafenib resistant cell line, identifying the acquisition of a point mutation in residue 842 in the second TKD (TKD2), resulting in the substitution of tyrosine (Y) for cysteine (C), thus generating a double mutant FLT3-ITD/Y842C receptor (Supplementary Table S2).

Treatment of these cells with type II TKIs sorafenib (Figure 1A) and quizartinib (Figure 1B), resistant cells showed an 80-fold (*p*=0.0003; area under the curve (AUC) 2-fold, *p*≤0.0001), and 881.5-fold (*p*=0.004; AUC 3.7-fold, *p*≤0.0001) increase in IC50 compared to FLT3-ITD sensitive cell lines, respectively (Supplementary Table S3). Consistent with previous reports of TKD2 mutations (22), these TKI resistant cells (FLT3-ITD/Y842C) showed increased sensitivity to type I TKIs midostaurin (IC50 0.50-fold, *p*=0.01; AUC 0.73-fold, *p*=0.01) (Figure 1C), and crenolanib (IC50 0.90-fold, *p*=0.04; AUC 0.85-fold, *p*=0.02) (Figure 1D) compared to the sensitive cells (Supplementary Table S3). No increase in Annexin V/PI staining was seen in TKI resistant cells treated with either sorafenib (mean viability = 88.2%) or quizartinib (mean viability = 85.9%). In contrast, treatment with the type II TKIs midostaurin (mean viability = 69.4%, p ≤ 0.0001) or crenolanib (mean viability = 38.2%, p ≤ 0.0001) promoted cell death. This effect was particularly notable in resistant cells compared to FLT3-ITD sensitive cell lines (p ≤ 0.0001) (Figure 1E-F; Supplementary Table S4).

**Figure 1.**
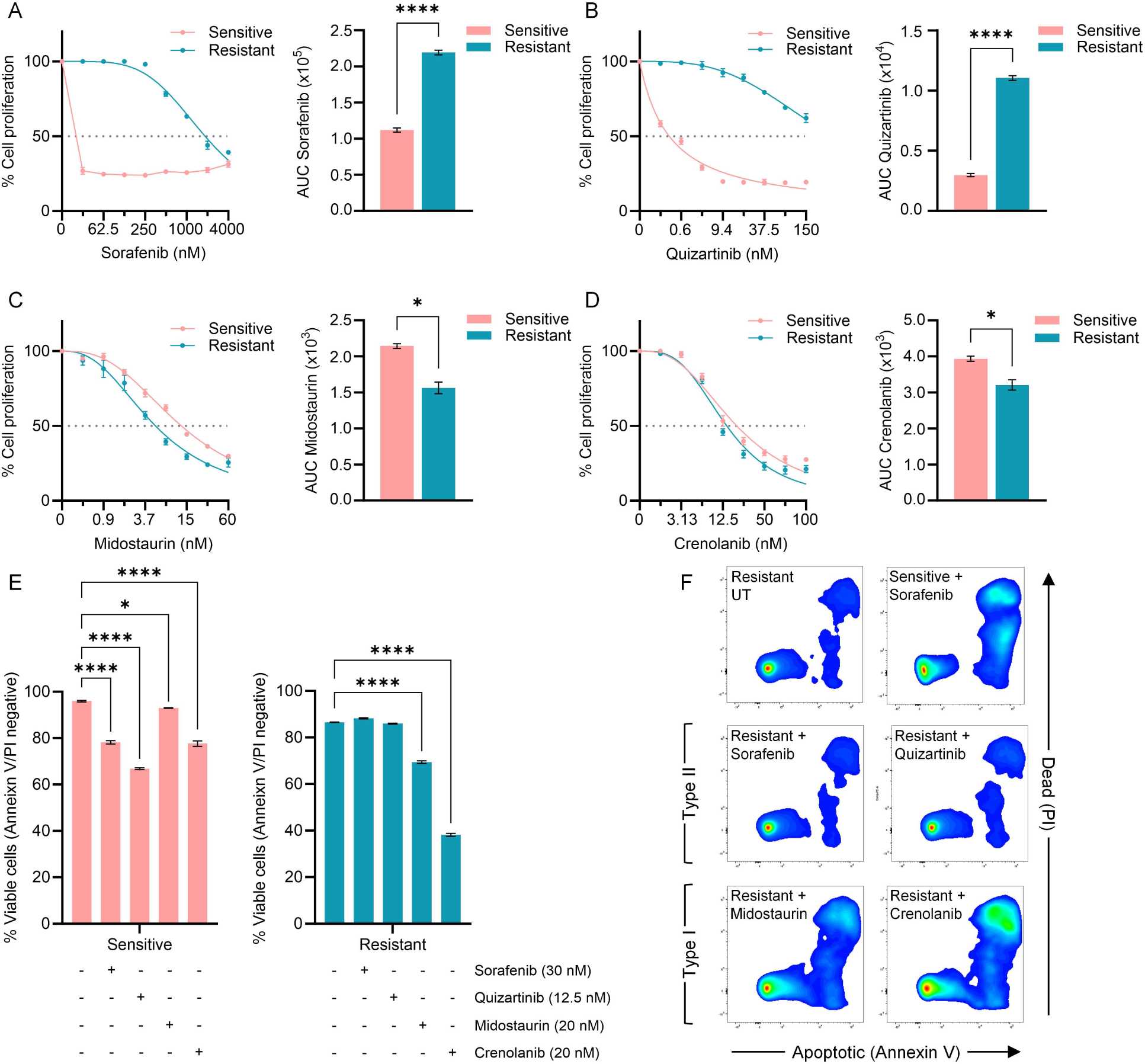
Comparison of tyrosine kinase inhibitor (TKI) sensitivity of MV4-11 FLT3-ITD cell lines. (A-D) Drug-response for FLT3-ITD sensitive and FLT3-ITD resistant cell lines was assessed using resazurin cell proliferation assays and area under the curve (AUC) was calculated and compared following treatment with type I and type II TKIs as single agents. Treatment with type II TKIs (A) sorafenib and (B) quizartinib confirmed FLT3-ITD adaptive resistance, whereas sensitivity was maintained following treatment with type I TKIs (C) midostaurin, and (D) crenolanib. Cell proliferation was assessed by resazurin fluorescence following 48-hour treatment (unpaired t-test with Welch’s correction; n=3 independent biological replicates). (E) Apoptosis and cell death was assessed via Annexin V/PI assay following 48-hour treatment with TKI’s as single agents in FLT3-ITD sensitive and FLT3-ITD resistant cell lines. Viable cells (Annexin V/PI negative) plotted as mean +/-SEM (ordinary one-way ANOVA; n=3 independent biological replicates). (F) Representative FACS image analysis indicating differences in cell death (PI) and apoptosis (Annexin V) after 48-hour treatment with Type I or II TKIs. Significance threshold of \**p*<0.05, \*\**p*<0.01 and \*\*\**p*<0.001

### Acquisition of FLT3-TKD2 mutations impact sorafenib and quizartinib binding

FLT3 is a 993 amino acid receptor tyrosine kinase with five immunoglobulin-like extracellular domains, a transmembrane domain, a cytoplasmic juxtamembrane (JM) domain, and two intracellular tyrosine kinase domains (TKD) (Figure 2A) (4). FLT3-ITD mutations disrupt the JM domain’s auto-inhibitory function, switching the receptor to its active conformation without FLT3L binding (31). To understand type II TKI resistance due to TKD2 point mutations, we performed *in silico* mutagenesis of the FLT3-TKD activation loop (aa 829 - aa 858) and analysed sorafenib and quizartinib binding (Figure 2B-E). Our resistance model includes ITD and a Y842C mutation in the second TKD (Supplementary Table S2). Both D835 and Y842 are within the activation loop and are inaccessible during auto-inhibitory conformation (Figure 2B-D) (9, 32). Type II TKIs (sorafenib and quizartinib) bind to the ATP-binding pocket of inactive FLT3 (yellow) (Figure 2B), while the active conformation sterically blocks binding due to phenylalanine 855 (F855) in the ‘back pocket’ (Figure 2D).

**Figure 2.**
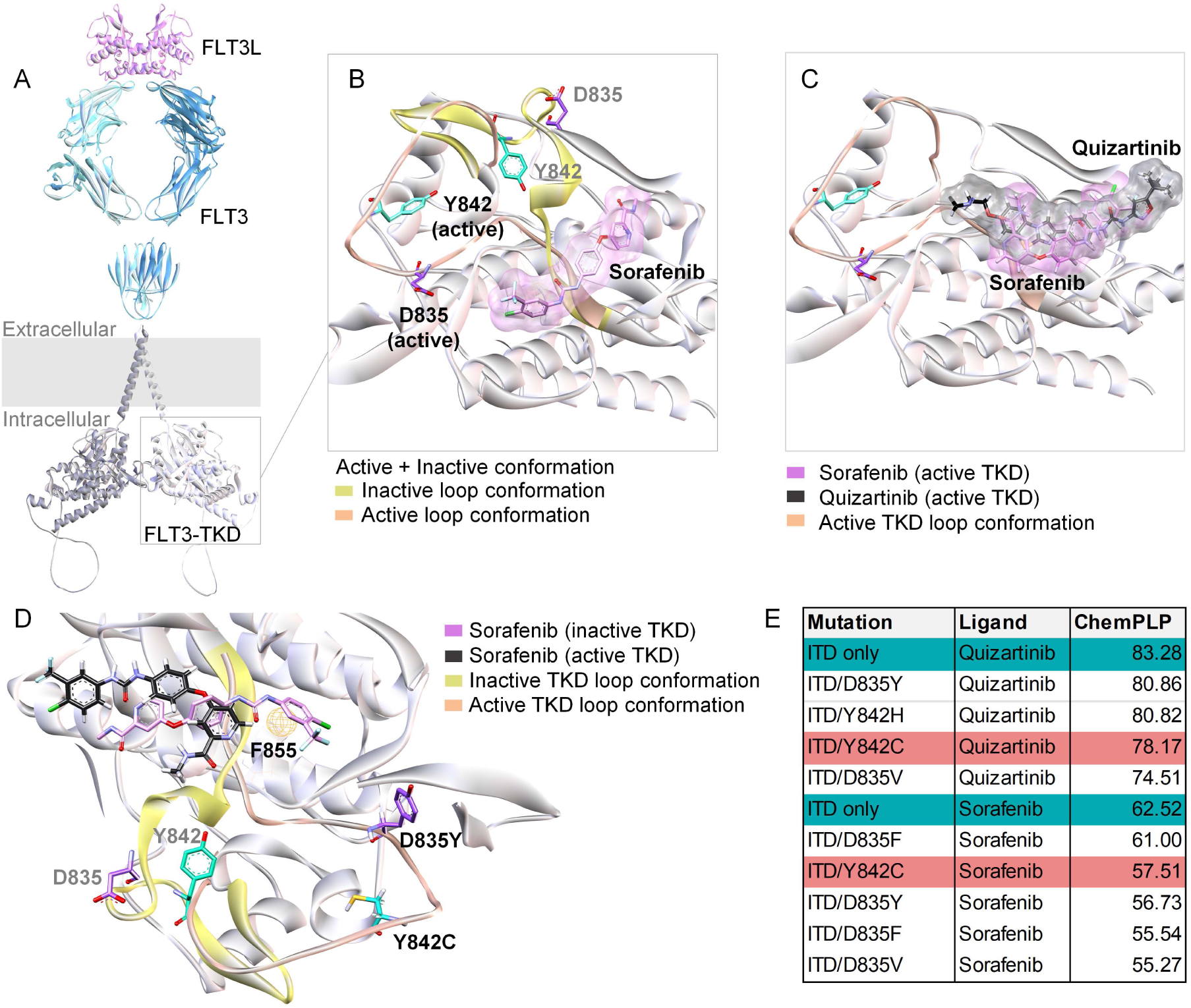
Structural features of active and inactive FLT3/TKD conformations and consequences for TKI binding. (A) A composite schematic of the FLT3 dimer bound to its ligand (FLT3L) (extracellular domain - PDB: 3QS9, transmembrane and TKD – AF-P36888-F1). (B) Structural overlay of the predicted active conformation for the FLT3 activation loop induced by the ITD mutation, highlighting differential locations of D835 (purple) and Y842 (green) residues with respect to the binding locations of sorafenib. Orange represents the active loop conformation whereas the inactive conformation is highlighted in yellow. (C) Active TKD confirmation demonstrating preferred docking poses of sorafenib and quizartinib. Both D835 (purple) and Y842 (green) residues, in the active confirmation, can influence the binding pocket. (D) Docking poses of sorafenib in both the active (black) and inactive (pink) FLT3-TKD conformations. The presence of D835 and Y842 mutations compromises the stability of the inactive TKD1 conformation, promoting activation. In the activated conformation, F855 (represented by the orange sphere) blocks the ability for sorafenib to anchor in its deep cleft (pink), causing it to bind superficially in the ATP pocket region (black). (E) Binding scores (ChemPLP) of type II TKI’s quizartinib and sorafenib to discreet double mutant FLT3 receptor models (higher score = better fit). Cell lines used throughout are indicated by green highlight (MV4-11 resistant) and pink highlight (MV4-11 sensitive).

Studies show that D835 and Y842 mutations cause loss of key hydrophobic and hydrogen bond interactions, destabilising the auto-inhibitory conformation and promoting the active loop, reducing drug affinity, especially with ITD mutations (9). ChemPLP binding affinity scores from *in silico* modelling (25) indicated reduced binding efficiency for quizartinib and sorafenib with TKD mutations. For FLT3-ITD/TKD-Y842C resistant cells, quizartinib binding decreased from 83.28 (ITD alone) to 78.17 (with Y842C), and sorafenib binding from 62.52 to 57.51 (Figure 2E). Thus, TKD point mutations decrease drug binding affinity by altering the activation loop conformation (Figure 2C-D).

### TKI resistant cells carry an increased dependency on DNA damage and repair pathways for survival

Differential signalling pathway analysis in resistance to type II TKIs, we performed by global, quantitative phosphoproteomics (n=3 independent biological replicates per cell line). Here we identified 1,469 unique phosphoproteins and 6,645 unique phosphorylated peptides (FDR 1%), with 1,335 phosphopeptides (20.1%) shown to be significantly altered in resistant cells to sensitive (*p*≤0.05). Two major clusters were shown to be differentially regulated in resistance (log2 ± 0.5, *p*≤0.05) (Figure 3A; Supplementary Table S5). Alterations in cell death and survival (*p*≤0.01), and DNA replication, recombination and repair (*p*≤0.009) were identified amongst the molecular and cellular functions modified by differential phosphorylation in both clusters (Figure 3B; Supplementary Table S6).

**Figure 3.**
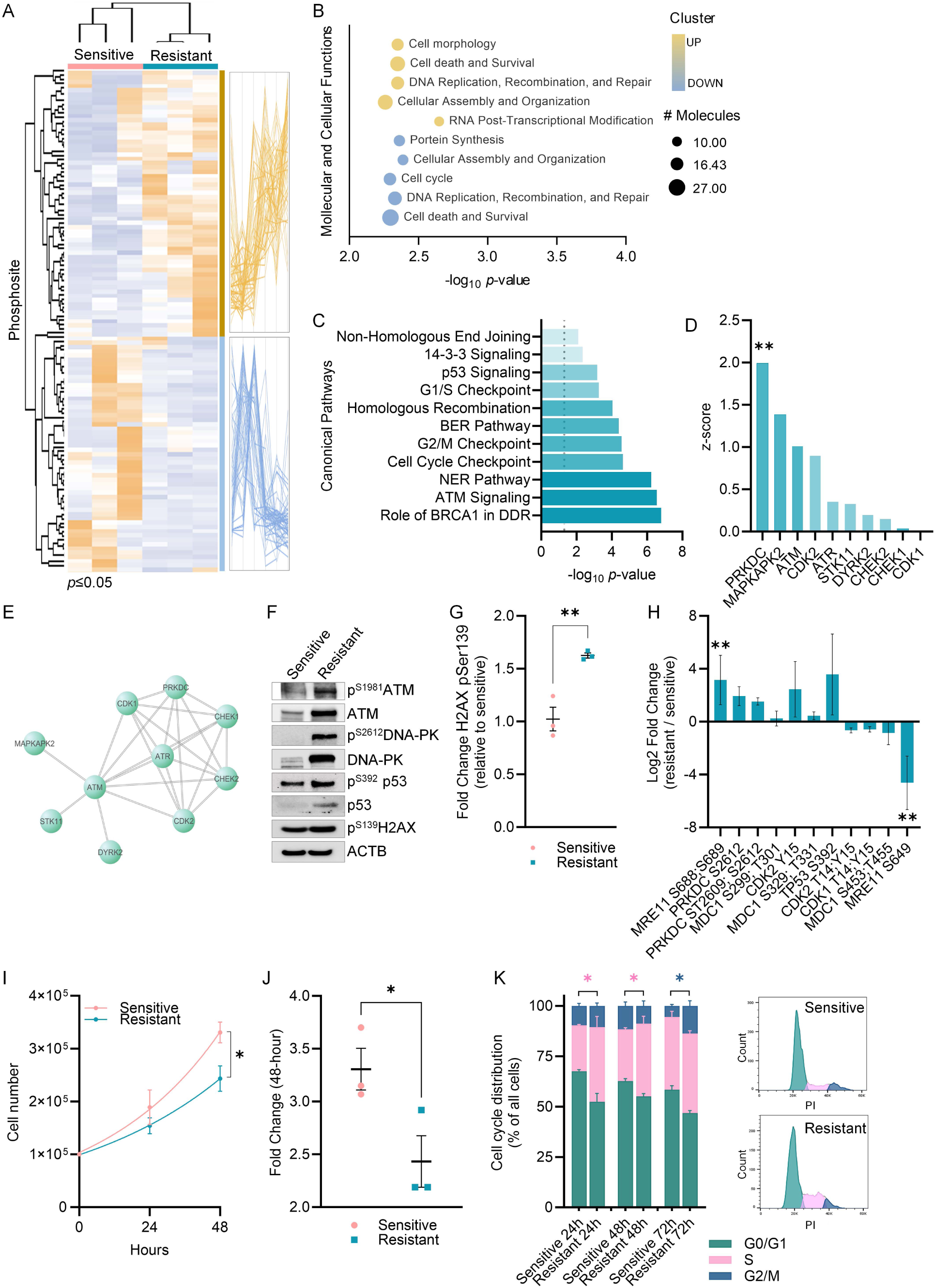
Phosphoproteomic analysis reveals overactivation of DNA damage and repair (DDR) signalling in FLT3-ITD resistant cells. (A) Identification of two independent phosphosite clusters in resistance. Heatmap (left) and cluster profile (right) of the precursor ion abundances for significantly up/down regulated phosphosites in three independent replicates (log2 ±0.5; *p*≤0.05). Yellow represents increased phosphorylation, whereas blue indicates phosphorylation is decreased in TKI resistant cells. (B) Molecular and cellular functions identified by Ingenuity Pathway Analysis (IPA) have been assigned to each cluster if significantly over-represented by phosphopeptides. (C) DNA damage and repair (DDR) canonical pathways identified by IPA as significantly associated with phosphorylation changes seen in FLT3-ITD resistant cells compared to sensitive cell lines. (D) Activity prediction for DDR kinases based on phosphorylation changes in substrates identified by Kinase-Substrate Enrichment Analysis (KSEA). (E) Protein-protein interaction network for DDR kinases identified by KSEA with a positive z-score in resistance (https://string-db.org/). (F) Western blot analysis further validated the increased expression and phosphorylation of DNA-PK (S2612), ATM (S1981), p53 (S392), and phospho-H2AX (S139) in resistance. (G) Phospho-H2AX (S139) western blotting data was quantified using Image Lab software and presented as a column graph comparing mean values ± SEM (n=3 independent biological replicates). Data was analysed by unpaired Student’s t-test. (H) Phosphorylation changes in key DDR signalling proteins identified by mass spectrometry. Values correspond to median log2 fold change in FLT3-ITD resistant cells compared to FLT3-ITD sensitive cell lines (log2 ±0.5; n=3 independent biological replicates). (I) Cell number based on relative cell count of FLT3-ITD sensitive (pink) and FLT3-ITD resistant (green) cell lines. Values at timepoints 0-, 24-, and 48-hours are shown (n=3 independent biological replicates). (J) Comparison of growth advantages of FLT3-ITD sensitive and FLT3-ITD resistant cell lines based on 48-hour fold change in cell density relative to day 0. Mean of triplicates ± SEM are shown. (K) Flow cytometry cell cycle analysis after staining with propidium iodide (PI). Cell cycle phase distribution shows the percentage of FLT3-ITD sensitive and FLT3-ITD resistant cells in the G0/G1, S, and G2/M phases of cell cycle at 24-, 48-, and 72-hour timepoints (n=3 independent biological replicates). Representative histogram of cell cycle profiles at 24-hours. Mean of triplicates ± SEM are shown. Significance threshold of \**p*<0.05, \*\**p*<0.01 and \*\*\**p*<0.001.

Indeed, constitutive DNA damage and repair (DDR) has previously been found to contribute to disease progression and therapeutic response in haematological malignancies, including FLT3-mutant AML (33). Analysis of the phosphorylation changes in resistant vs sensitive cells in DDR proteins identified BRCA1 (*log10 p*=6.82), ATM (*log10 p*=6.57), and Nucleotide excision repair (NER) (*log10 p*=6.26) pathways as the top 3 DDR-associated signalling pathways in resistance (Figure 3C; Supplementary Table S7) corroborating studies of resistance signalling in FLT3-ITD/D835V/Y mutations (26). Accordingly, Kinase-Substrate Enrichment Analysis (KSEA) predicted the key DDR kinases, DNA-dependent Protein Kinase PRKDC (DNA-PK; *z-score* = 1.99, *p*=0.02), and ATM kinase (*z-score*= 1.01), to be increased in activity in TKI resistant cells with a positive *z-score* indicating activation (Figure 3D; Supplementary Table S8). Protein-protein interaction analysis of DDR kinases identified by KSEA with a positive z-score (indicating activation) revealed four separate nodes of kinase interaction, predominantly connected through the ATM kinase (Figure 3E), validated by immunoblotting (Figure 3F). Increase in phosphorylation of the key DNA damage marker H2AX (S139) was seen in resistant cells (*p*=0.006) (Figures 3F-G), so we analysed the phosphoprotein changes in key proteins regulating ATM-driven DDR signalling (log2 ± 0.5) (Figure 3H, Supplementary Table S9), including the Double-Strand Break (DSB) repair protein MRE11 (S688; S689; *p*=0.03), whereas its DNA repair inhibitory site (S649) was significantly decreased in phosphorylation (*p*=0.01). Increased phosphorylation of the activating sites of DNA-PK, S2612 (log2 fold=2.4) and S2609 (log2 fold=1.3), as well as cellular tumour antigen p53, S392 (log2 fold=0.5), were also identified via phosphoproteomics (Figure 3H) and confirmed via immunoblotting (Figure 3F).

### TKI resistant cells show decreased cell growth and proliferation

Alterations in cell cycle regulation were significantly overrepresented in the cluster with decreased phosphorylation in resistant cells (Figure 3B; Supplementary Table S6). IPA analysis predicted significant associations between phosphorylation changes in TKI-resistant cells and G2/M (log10 p=4.58) and G1/S (log10 p=3.29) cell cycle checkpoint regulation (Figure 3C; Supplementary Table S7). Given that cell cycle checkpoint activation controls DDR response (34), we assessed growth profiles of FLT3-ITD sensitive and resistant cell lines using cell proliferation (Figure 3I-J) and cell cycle (Figure 3K) assays.

Resistant cells displayed a 1.36-fold decrease in cell proliferation at the 48-hour timepoint compared to sensitive cells (p=0.02) (Figure 3I-J). Flow cytometry analysis showed differences in all cell cycle phases between sensitive and resistant lines, with a higher percentage of resistant cells in the S phase at 24 and 48 hours (p=0.01), and in the G2/M phase at 72 hours (p=0.03) (Figure 3K; Supplementary Table S10).

### High ATM expression associated with worse overall outcomes in in paediatric AML and FLT3-mutant adult AML

To assess the clinical relevance of ATM expression in AML, we examined ATM expression in adult FLT3-ITD mutant patients at diagnosis and paediatric patients at diagnosis, progression, and relapse, using data from the Beat AML and Therapeutically Applicable Research to Generate Effective Treatments (TARGET) databases (33,34). In FLT3-ITD AML patients, high ATM expression is associated with shorter median overall survival (OS) compared to low ATM expression (n=68; *p*=0.019; 95% CI=1.1886 log-rank) (Figure 4A). Among paediatric patients, high ATM expression is linked to shorter event-free survival (n=239; *p*=0.012; 95% CI=1.1489 log-rank) (Figure 4B), shorter OS (*p*=0.013, 95% CI=1.565 log-rank) (Figure 4C), and shorter OS at relapse (n=125, *p*=0.048, 95% CI=1.552 log-rank) (Figure 4D). Increased ATM expression was observed at relapse compared to diagnosis (n=242, *p*=0.0015, Two-Tailed Welsch T-Test) (Figure 4E).

**Figure 4.**
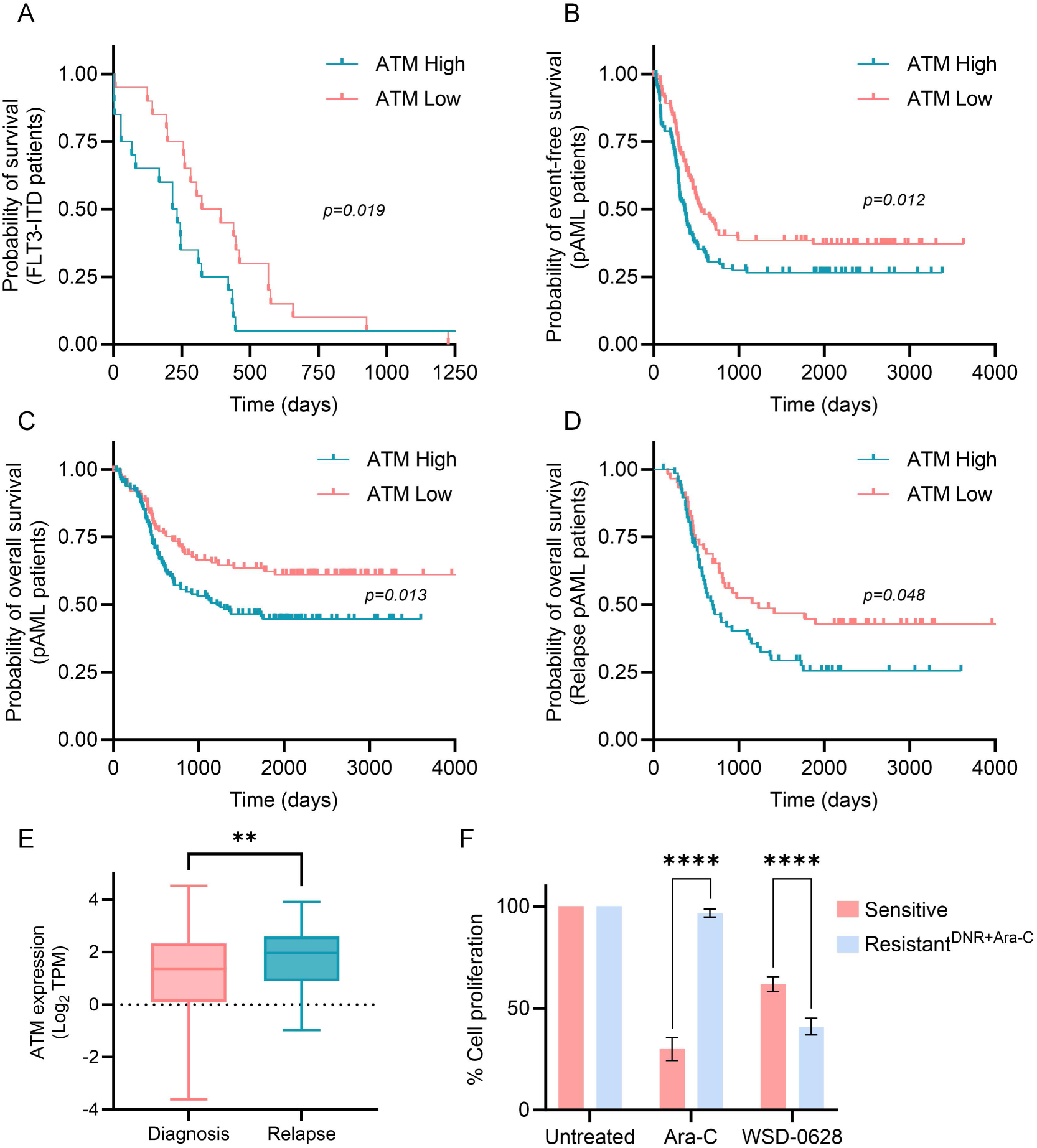
Analysis of publicly available survival and expression data stratifies patients based on ATM expression. (A) Patient data from BEAT AML Vizome database were downloaded and expression values (TPM) as well as clinical information were obtained for 68 FLT3-ITD adult AML patients. Patients were separated into high and low ATM expression as determined by TPM value where high expression referred to the top 25% of patients and low expression the bottom 25% of patients (Q1/Q4 split). Survival analysis was then performed using the Kaplan-Meier model and the Log-rank, Mantel-Cox statistical test used to compare overall survival at diagnosis. Next, the Therapeutically Applicable Research to Generate Effective Treatments (TARGET) initiative, phs000465, was downloaded and again expression values (TPM) as well as clinical information were obtained for 285 paediatric AML patients. (B) Event free survival and (C) overall survival in all paediatric AML (pAML) cases, and (D) overall survival in the relapse setting. (E) ATM expression (Log2 TPM) was then compared across all AML patients from the TARGET database segregated by diagnosis or relapse subtype. Students t-test was performed for statistical comparison. (F) Resazurin proliferation (percentage compared to untreated) assays of FLT3-ITD sensitive and FLT3-ITD DNR (daunorubicin) and Ara-C (cytarabine) resistant cell lines after 48-hour exposure to 25 µM AraC or 250 nM WSD-0628 (minimum of n=3 independent biological replicates). Significance threshold of \**p*<0.05, \*\**p*<0.01 and \*\*\**p*<0.001 (Two-Way ANOVA).

To evaluate the sensitivity of paediatric AML cells treated with the clinically relevant brain-penetrant ATM inhibitor WSD-0628, we exposed standard-of-care sensitive (SOC) sensitive and resistant AML cells (cytarabine and daunorubicin) for 48 h and measured viability. Both SOC sensitive and resistant cells demonstrated high sensitivity to WSD-0628, however, SOC-resistant cells were significantly more sensitive than SOC-sensitive cells (*p*≤0.0001, Two-Way ANOVA) (Figure 4F).

### Type II TKI resistant cells reside in a state of high-level oxidative stress and oxidative DNA damage

Activation of DDR pathways, including ATM signalling, is triggered by the presence of DNA DSBs (35) commonly caused by excessive levels of ROS, considered a driver of disease progression in FLT3-mutant AML (29, 36, 37). Additionally, FLT3 inhibition itself is reported to result in the accumulation of ROS, consequently activating ATM signalling to maintain redox homeostasis (38). Commensurate with SOC resistant AML cells and patients (36, 37, 39), TKI-resistant cells showed increased cytoplasmic superoxide (DHE positive fluorescence) compared to sensitive cells, with a 1.93-fold increase in ROS levels (*p*=0.004) (Figure 5A-B). We next tested cell proliferation following treatment with the ROS scavenger N-Acetylcysteine (NAC, 1.25–20 mM). TKI-resistant cell lines showed significantly higher proliferation rates in 20 mM NAC (*p*=0.03) (Figure 5C). Phosphoproteomic analysis of ROS-associated canonical pathways in resistant cells identified significant increases in the ERK/MAPK (log10 *p*=8.9), PI3K/AKT (log10 *p*=3.5), and NRF2-mediated oxidative stress response (log10 *p*=1.45) pathways (Figure 5D).

**Figure 5.**
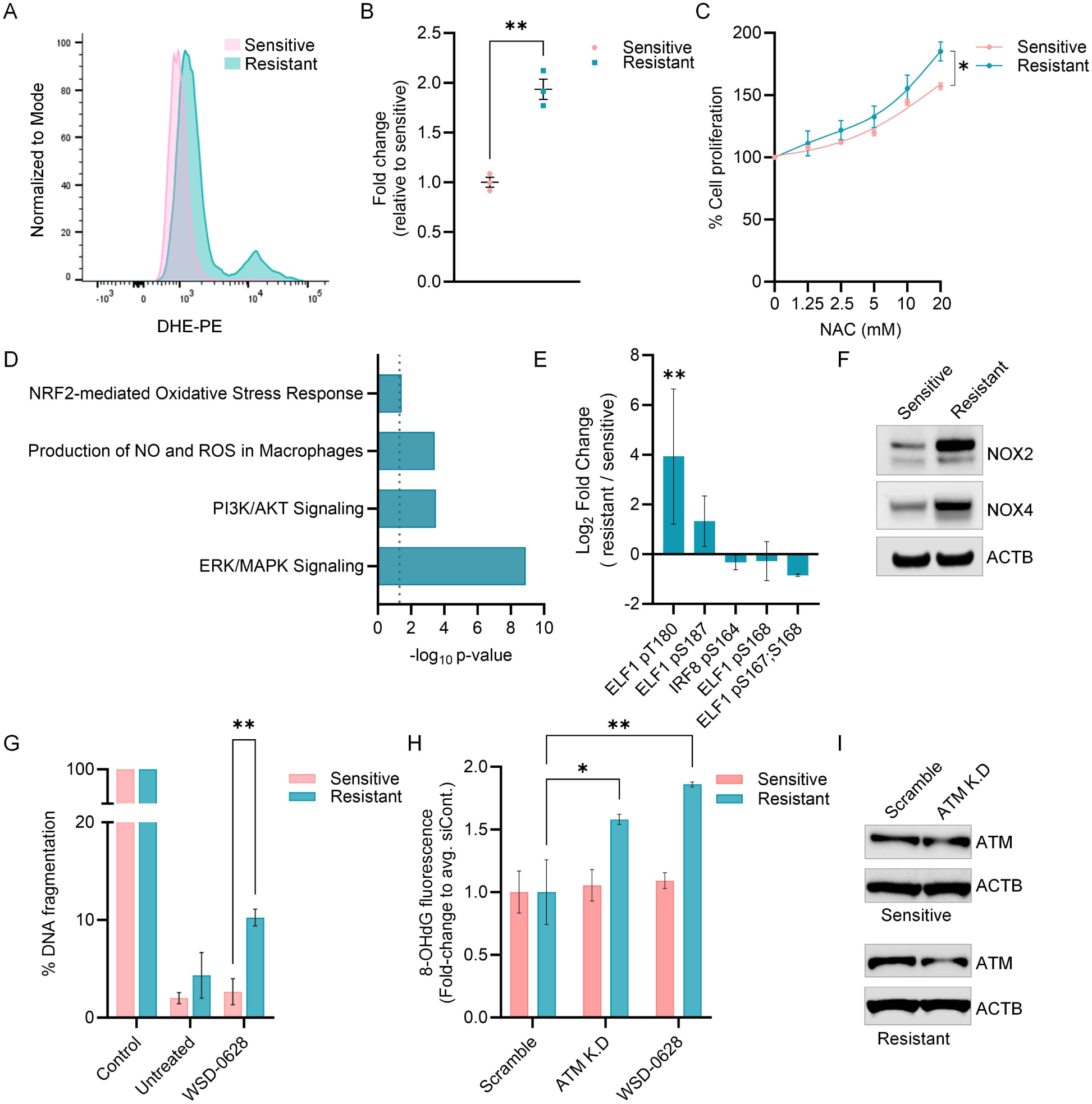
FLT3-ITD resistant cells display higher levels of reactive oxygen species (ROS) in comparison to FLT3-ITD sensitive cell lines. (A) Flow cytometry histogram overlay of cytoplasmic ROS (superoxide) levels in FLT3-ITD sensitive (pink), and FLT3-ITD resistant (green) cell lines. ROS levels were assessed by DHE-PE fluorescence and analysed using FlowJo software. (B) Geometric means were used for ROS quantification and fold change comparison, presented as a column graph as mean values ± SEM (n=3 independent biological replicates). (C) Analysis of cell proliferation for FLT3-ITD sensitive and FLT3-ITD resistant cells in increasing concentrations of ROS scavenger NAC. Cell proliferation was assessed by resazurin assay following 48-hour treatment (n=3 independent biological replicates). Mean of triplicates ± SEM are shown. (D) ROS-associated canonical pathways identified by Ingenuity Pathway Analysis (IPA) as significantly associated with phosphorylation changes seen in FLT3-ITD resistant cells compared to FLT3-ITD sensitive cell lines. (E) Phosphorylation changes in NOX2 transcription factors ELF1 and IRF8 in FLT3-ITD resistant cells compared to FLT3-ITD sensitive. (F) Western blotting reveals increased protein expression of NADPH oxidase isoforms NOX2 and NOX4 in resistance. (G) ICC quantification of DNA fragmentation (TUNEL-positive) in FLT3-ITD sensitive and FLT3-ITD resistant cells in untreated conditions and after 48-hours treatment with 250 nM of the ATM inhibitor WSD-0628. (H) ICC quantification of oxidative DNA damage (8-OHdG-positive) in FLT3-ITD sensitive and FLT3-ITD resistant cells carrying knockdown (K.D) of ATM, scramble control or treated for 48-hour with 250 nM WSD-0628. Significance threshold of \**p*<0.05, \*\**p*<0.01 and \*\*\**p*<0.001 (n=3 independent biological replicates).

Based on these data, we assessed whether the increased levels of ROS could result from differences in the expression of the NADPH oxidases (NOX2/4). Indeed, phosphoproteomic analysis identified ELF1 and IRF8 transcription factors as differentially phosphorylated in resistance, responsible for the transcriptional regulation of NOX2 and its associated subunits (Figure 5E) (40). Consistent with these findings, NOX2 and NOX4 protein expression showed increased expression in TKI resistant cells (Figure 5F). Finally, assessment of DNA damage in resistant cells revealed similar levels of DNA fragmentation in FLT3-ITD sensitive and TKI resistant cells under normal conditions. However, blocking DNA repair via inhibition of ATM using WSD-0628, resulted in a significant increase in DNA fragmentation in TKI resistant cells (3.8-fold, *p*=0.008) (Figure 5G). Similarly, blocking DNA repair through the pharmacological inhibition of ATM led to a significant increase in oxidative DNA damage in resistance, indicated by the presence of the oxidized DNA nucleoside guanosine (8-OHdG) (Figure 5H). Equally, molecular inhibition of ATM (Figure 5I) led to a significant increase in oxidative DNA damage, analogous to that seen with WSD-0628 treatment, restricted to TKI resistant cells (Figure 5H).

### ATM inhibition reduced cell proliferation in vitro and increased survival in vivo

To assess the therapeutic potential of targeting DDR signalling following the development of adaptive resistance to type II TKIs, we performed cytotoxicity analysis of FLT3-ITD sensitive and TKI resistant cells using the DNA-PK inhibitor peposertib (formerly known as M3814), ATM inhibitor KU-60019, and a second, more potent brain penetrant ATM inhibitor, WSD-0628, as single agents (Supplementary Figure S3A-C). Both FLT3-ITD sensitive and TKI resistant cell lines responded to DNA repair inhibition to all three single agents. Treatment with peposertib (Supplementary Figure S3A) and KU-60019 (Supplementary Figure S3B) did not show significant differences in sensitivity between FLT3-ITD sensitive and TKI resistant cells, even in the micromolar dose range. However, Both TKI sensitive and resistant cells showed nanomolar sensitivity to WSD-0628, with TKI resistant cells significantly more sensitive (IC50 116.5 nM TKI resistant; IC50 183.1 nM FLT3-ITD sensitive; AUC resistant vs sensitive 0.60-fold, *p*=0.003) (Figure 6A-C, Supplementary Figure S3C). We assessed the response of TKI resistant cells to WSD-0628 alone, and in combination with the type II TKIs sorafenib or quizartinib (Figure 6A-C, Supplementary Figure S3D, E). The combination of sorafenib with WSD-0628 (sorafenib 31.2 nM + WSD-0628 125 nM) significantly decreased cell proliferation by 1.8-fold (*p*=0.0008) compared to treatment with sorafenib alone (Figure 6A, Supplementary Figure S3D). A similar response was seen for quizartinib (quizartinib 12.5 nM + WSD-0628 125 nM), with a 1.6-fold reduction (*p*≤0.0001) (Figure 6B, Supplementary Figure S3E). Drug combinations were then assessed for synergistic interactions via the computational synergy analysis BLISS, however, both combinations only showed additive effects (Bliss score WSD-0628 + sorafenib: 4.09; Bliss score WSD-0628 + quizartinib: 4.68) (Supplementary Figure S3F, S3G, respectively) (41)

**Figure 6.**
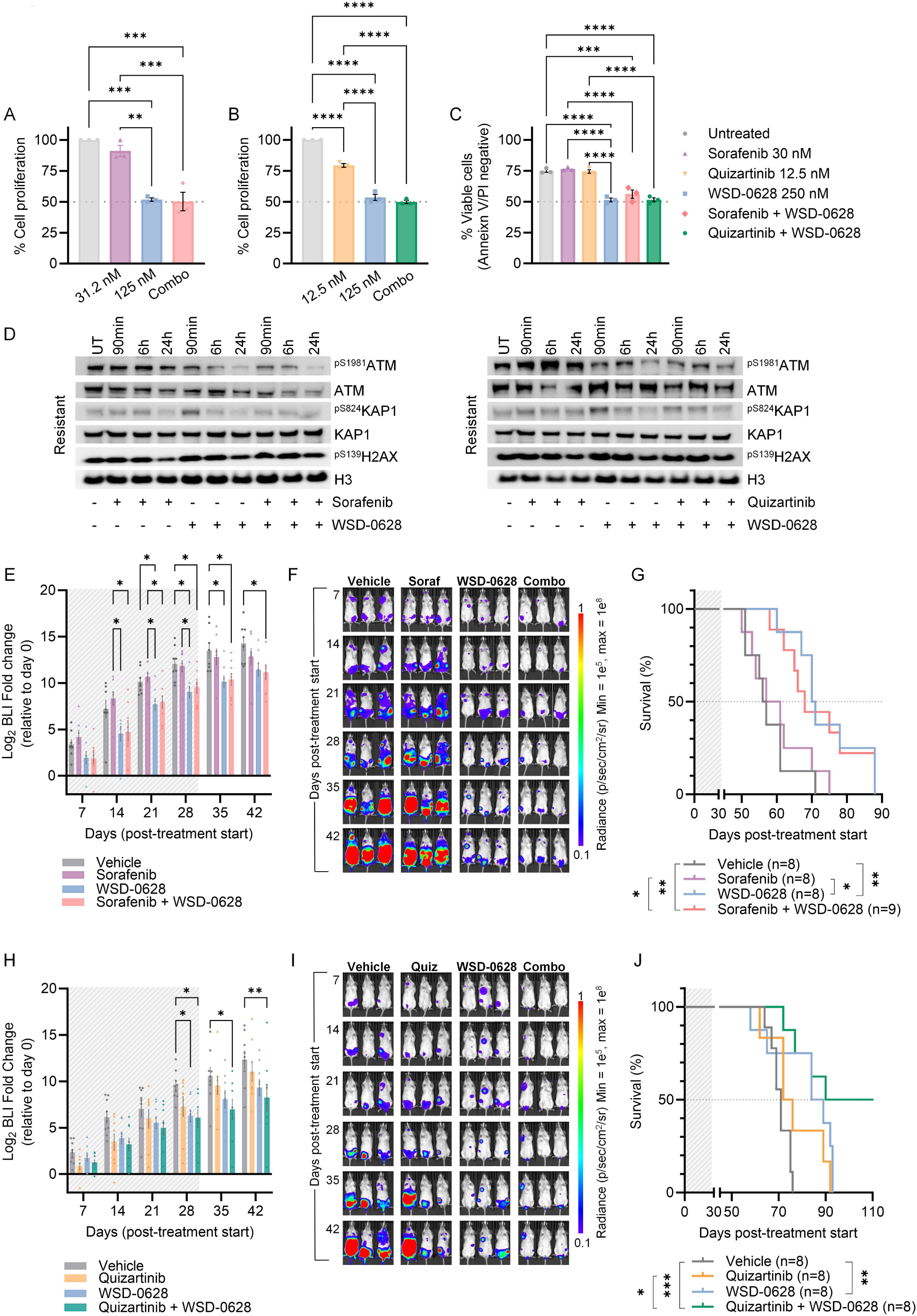
Sensitivity to ATM inhibition alone and in combination with FLT3 inhibitors. Resazurin proliferation (percentage compared to untreated) of FLT3-ITD resistant cell lines after 48-hour exposure to WSD-0628 alone, and in combination with FLT3 inhibitors (A) sorafenib and (B) quizartinib. Values shown as mean ± SEM (n=3 independent biological replicates). (C) Annexin V/PI apoptosis assay following 48-hour exposure to TKI as single agents or in combination with WSD-0628 in FLT3-ITD resistant cell lines. Values presented as mean ± SEM (ordinary one-way ANOVA; n=3 independent replicates). (D) Phosphorylation changes in proteins regulating the activation of DNA damage and repair following treatment with ATM inhibitor alone, and in combination with sorafenib, or quizartinib, measured by Western blotting. (E-J) MV4-11 FLT3-ITD resistant cell lines were injected into the lateral tail vein of NOD-Rag1null IL2rgnull (NRG) mice. Treatment commenced once BLI reached a mean radiance of 1 × 10^6^ p/s. Sorafenib, quizartinib, and WSD-0628 were administered once daily for 4 weeks. (E-F) *In vivo* monitoring of leukemia burden using bioluminescence BLI imaging over time (representative BLI images presented, shaded area indicates treatment time) of mice treated with WSD-0628, sorafenib or the combination. (G) Kaplan-Meier survival analysis of MV4-11–Luc+ FLT3-ITD resistant cells (n= 8 mice per group, shading indicating treatment duration) treated with WSD-0628, sorafenib, or combination (Log-rank, Mantel-Cox). (H-I) Monitoring of leukemia burden in the second study using bioluminescence BLI imaging over time (representative BLI images presented, shaded area indicates treatment time) of mice treated with WSD-0628, quizartinib or the combination. (J) Kaplan-Meier survival analysis of MV4-11–Luc+ FLT3-ITD resistant cells (n= 8 mice per group, shading indicating treatment duration) treated with WSD-0628, quizartinib, or combination (Log-rank, Mantel-Cox). Significance threshold of \**p*<0.05, \*\**p*<0.01 and \*\*\**p*<0.001.

(Figure 6A-B). Consistently, treatment with WSD-0628 alone and in combination with sorafenib, or quizartinib, promoted significant levels of cell death compared to the untreated controls and either sorafenib or quizartinib alone (mean viability = 56.1%, *p*=0.0001 and 51.4%, *p*≤0.0001 respectively) (Figure 6C, Supplementary Table S11).

To evaluate the downstream signaling response to WSD-0628 alone or in combination, changes in ATM kinase and associated DNA damage and repair proteins were assessed via immunoblotting. ATM auto-phosphorylates at S1981 in response to DNA damage (35), and promotes chromosome relaxation by phosphorylating KAP1 at S824 (42). Phosphorylation induces KAP1 co-localisation with γH2AX at damage sites, facilitating homologous recombination (HR) and non-homologous end joining (NHEJ) repair (42, 43). Treatment with WSD-0628 alone or combined with TKIs decreased phosphorylation of ATM (S1981) and H2AX (S139) in TKI-resistant cells, while TKI treatment alone did not (Figure 6D).

To evaluate the anti-AML potential of ATM inhibition, we engrafted NOD-*Rag1^null^ IL2rg^null^* (NRG) mice with luciferase transduced MV4-11 FLT3-ITD model of adaptive resistance to type II TKIs. Mice were randomised on detection of BLI and treated with vehicle, WSD-0628, sorafenib, quizartinib, or WSD-0628 combined with either sorafenib or quizartinib (Figure 6E-J). After 4 weeks, leukaemia burden (BLI) significantly decreased in mice treated with WSD-0628 alone (*p*=0.02) and combined with sorafenib (*p*=0.04), but not with sorafenib alone (Figure 6E-F). Sorafenib monotherapy did not extend survival compared to the vehicle group (59 days vs. 56.5 days) (Figure 6G). Mice treated with WSD-0628 monotherapy survived longer than both the vehicle group (70.5 days vs. 56.5 days, *p*=0.006) and the sorafenib group (70.5 days vs. 59 days, *p*=0.02). WSD-0628 combined with sorafenib significantly extended survival compared to the vehicle (68 days vs. 56.5 days, *p*=0.003) and sorafenib groups (68 days vs. 59 days, *p*=0.03), but not compared to WSD-0628 monotherapy (68 days vs. 70.5 days) (Figure 6G).

Next, we evaluated the effects of WSD-0628 and quizartinib as monotherapies, and in combination. After 4 weeks, mice treated with WSD-0628 alone (*p*=0.03) and in combination with quizartinib (*p*=0.01) showed a significant reduction in leukaemia burden, whereas those treated with quizartinib alone did not (Figure 6H-I). Quizartinib monotherapy did not extend survival compared to controls (74 vs. 71 days). In contrast, WSD-0628 alone significantly extended survival (86.5 vs. 71 days, *p*=0.009). The combination of WSD-0628 and quizartinib further extended survival compared to vehicle- (90 vs. 71 days, *p*=0.0003) and quizartinib-treated groups (90 vs. 74 days, *p*=0.03), but not significantly compared to WSD-0628 alone (90 vs. 86.5 days) (Figure 6J). After 120 days, 50% of mice treated with the combination showed no signs of leukaemia (Figure 6J).

To assess if ATM inhibition promotes similar responses in *in vivo* models of FLT3-ITD TKI sensitive models MV4-11 FLT3-ITD cell lines were engrafted and treated with vehicle, WSD-0628, or sorafenib (Supplementary Figure S4). After 4 weeks, sorafenib alone significantly reduce leukaemia burden (*p*=0.0167) compared to vehicle controls. WSD-0628 did not significantly reduce BLI radiance over 4 weeks but stalled leukaemia progression, showing a significant decrease after 5 weeks (*p*=0.0347) (Supplementary Figure S4A). Sorafenib significantly extended survival (53 vs. 40 days, *p*=0.0155), as did WSD-0628 monotherapy (55 vs. 40 days, *p*=0.0006) (Supplementary Figure S4B).

## DISCUSSION

The development and approval of TKIs targeting FLT3 have improved treatment strategies for FLT3-ITD AML patients. Currently, two TKIs, midostaurin and quizartinib, are FDA-approved for use with induction chemotherapy in newly diagnosed FLT3-mutant AML. Additionally, sorafenib is often used off-label post allo-HCT or following resistance in relapsed FLT3-ITD AML patients who have also received gilteritinib salvage treatment (16). For the treatment of pAML, sorafenib is the most extensively studied first-generation FLT3 inhibitor (44), with reports showing it can be safely combined with standard of care chemotherapy to improve outcomes in high allelic ratio (HAR) (AR > 0.4) FLT3-ITD pAML (AAML1031) (17).

However, relapse following TKIs remains a challenge, and the lack of alternative treatments for patients who only transiently respond to current FLT3-targeted therapies contributes to low survival rates. Here, we present a comprehensive phosphoproteomic analysis of FLT3-ITD AML resistant to type II TKIs, sorafenib and quizartinib. We found that ROS-driven DDR signaling, particularly through ATM regulation, is overactivated in TKI-resistant cells, promoting survival despite treatment. This study identifies key factors controlling TKI-resistant cell survival, providing crucial information for designing patient-specific therapies targeting both common and divergent oncogenic signaling pathways.

Our phosphoproteomics analysis revealed that ATM-driven DNA repair signalling is crucial for cell survival and therapy resistance in FLT3-ITD/TKD cells. The DDR pathway, activated by endogenous DNA damage often caused by ROS, is a key factor. High ROS levels drive progression in FLT3-ITD AML (29, 36, 37). We confirmed that TKI-resistant cells produce more ROS than TKI-sensitive cells due to increased NOX2/4 protein expression, indicating a redox imbalance that aids survival. This pattern is also seen in AML patients resistant to SOC (37). Reducing ROS increased the proliferation of resistant cells, suggesting less reliance on DDR response and more energy for proliferation.

Acute FLT3 inhibition induces ROS accumulation, activating ATM signalling to maintain redox homeostasis (48). We observed increased phosphorylation of histone H2AX (γH2AX pSer139), ATM (S1981), and DNA-PK (S2609, S2612) kinases in resistant cells, highlighting their role in DDR signalling regulation (45). Inhibiting ATM in TKI-resistant cells significantly increased DNA fragmentation and oxidative DNA damage, which did not occur in TKI-sensitive cells. This underscores the importance of enhanced DNA repair via ATM for cell survival under oxidative stress in TKI-resistance, presenting a novel therapeutic vulnerability in TKI resistance and as a salvage therapy post-TKI failure.

In the current clinical management of leukaemia, minimal residual disease (MRD) detection provides critical insight into the remission status of patients and has significantly contributed to the overall improvement in survival rates (46). However, pAML patients with higher MRD-positive rates after standard induction therapies are at an increased risk of relapse and have worse overall survival (47). Importantly, we identified that pAML patients with high-level expression of ATM fare significantly worse across all disease settings than those with low ATM expression.

Increased activity and expression of NOX enzymes responsible for high oxidative stress in high-risk AML are well-documented, but therapeutic targeting remains challenging (37, 48). Our findings suggest that ROS-mediated ATM signalling drives a constitutive DDR feedback loop that sustains cell survival under ROS-induced stress (38, 48), presenting an exciting therapeutic opportunity. Given the interest in ATM kinase inhibitors across various cancers (49), we evaluated ATM inhibition as a strategy against SOC- and FLT3-ITD/TKD-resistance. Treatment with the clinically relevant ATM inhibitor WSD-0628, alone and in combination, successfully reduced resistant cell proliferation *in vitro* and decreased phosphorylation of ATM kinase and γH2AX, critical for DSB repair fidelity (43). *In vivo*, ATM inhibition reduced leukaemia burden and significantly increased survival of mice engrafted with TKI-resistant cells, either as a single agent or in combination with TKIs sorafenib and quizartinib.

Clinically, WSD-0628 is being tested with radiation therapy for glioblastoma (NCT05917145), and new studies are underway for paediatric diffuse midline gliomas. Our data suggest that ATM inhibition could be a valuable addition to standard induction therapies, used in consolidation, or as a salvage therapy at relapse in R/R FLT3-mutant AML. This approach may address the high MRD-positive rates and poor outcomes associated with high ATM expression, providing a novel and promising therapeutic strategy for improving patient survival.

## ACKNOWLEDGMENTS

N. Smith from the University of Newcastle Analytical and Biomolecular Research Facility (ABRF) provided MS support. The Academic and Research Computing Support (ARCS) team, within Digital Technology Solutions at the University of Newcastle, provided high-performance computing (HPC) infrastructure supporting bioinformatic analyses.

The ATM inhibitor, WSD-0628, used in this study was supplied by Wayshine Biopharm International Ltd, Corona, California under a materials transfer agreement with the University of Newcastle, Callaghan, Australia.

## AUTHOR CONTRIBUTIONS

D.E.S., Z.P.G., A.M., T.M. and M.D.D. conceived and designed the study and interpreted the results; D.E.S., Z.P.G., A.M., T.M., L.C., G.D.I., and M.D.D. conducted the experiments and performed data analysis; B.C.T., H.P.M., H.C.M., R.J.D., T.B., M.L.P., I.J.F., E.R.J., N.D.S., D.A.S., B.N., and N.V. helped with experimental work and/or interpretation of results; P.C., assisted with bioinformatic analyses; D.M., W.J., A.K.E., J.R.S., J.C., F.A. provided discipline-specific expertise; J.C., F.A., J.R.S., A.K.E., and A.H.W. assisted with obtaining and processing of patient samples; D.E.S., Z.P.G., A.M.D., and M.D.D. wrote and edited the manuscript; and all authors discussed the results and commented on the manuscript.

## Data and materials availability

MTA agreements and publicly accessible data as described in Materials and Methods.

## DATA SHARING STATEMENT

The mass spectrometry proteomics data has been deposited to the ProteomeXchange Consortium (http://proteomecentral.proteomexchange.org) via the PRIDE partner repository with the dataset identifier PXD053329 and 10.6019/PXD053329

## MANUSCRIPT FEATURES

Manuscript - Abstract: 250 words, main text: 4020 words, 6 Figures. Supplementary Information - Supplementary Materials and Methods, 4 Supplementary Figures. Supplementary Excel file - 11 Supplementary Tables.

