## Supplementary Information for "Adaptive resistance to FLT3 inhibitors is potentiated by ROS-driven DNA repair signalling"

### SUPPLEMENTARY METHODS

*Cell culture*

Three human MV4-11 AML cell lines (FLT3-ITD sensitive, TKI resistant and SOC resistant) were maintained in standard culture conditions (5% $CO_2$, 37˚C) in DMEM medium (Thermo Fisher Scientific, Waltham, MA, USA) with the addition of 10% FBS, and 20 mM HEPES (N-2-hydroxyethylpiperazine-N'-2-ethanesulfonic acid). FLT3-mutant lines are factor-independent and were therefore maintained in growth factor-free media. All cell lines were routinely assessed using a MycoAlert mycoplasma detection kit and confirmed to be free of mycoplasma contamination (Lonza; Basel, Switzerland).

*In vitro development of adaptive resistance*

TKI resistant MV4-11 cells were developed through serial passaging of cells in increasing doses of sorafenib (Selleckchem, Houston, TX, USA) from 2.5 nM-1280 nM over 20 weeks. Cell viability and relative growth rates were assessed via Trypan Blue exclusion as previously described (1) and were considered resistant when 50% viability was maintained in the presence of drug (Supplementary Figure S1). Adaptive resistance was validated via next generation sequencing (NGS) as below, and additional mutations identified are listed in Supplementary Table S2. The effect of TKIs on cellular growth and proliferation of MV4-11 cells was determined using resazurin growth and proliferation assay as previously described (2). Cell death was measured by Annexin V apoptosis assay as previously described (3). Cell cycle analysis was performed using propidium iodide flow cytometry as previously described (2).

To make SOC resistant MV4-11 cells were serial passaged with increasing doses of cytarabine over a 12-month period, by which cells were then able to successfully proliferate whilst in the presence of 500μM of cytarabine. These cytarabine resistant MV4-11 cells were then serially passaged with increasing doses of daunorubicin to reach a maximum dose of 12 nM. Resistance was confirmed via cytotoxicity assays and cell colony forming assays. Long term maintenance of resistance has been confirmed without the need for reselection via cytarabine. Resistant MV4-11 cells have been passaged for a further 12 months without re exposure to cytarabine, followed by cytotoxicity assays to demonstrate the cells resistance phenotype was maintained.

*Next generation sequencing (NGS)*

DNA was extracted using a Wizard SV Genomic DNA Purification System (Promega, Madison, WI, USA), as per manufacturer's instructions. Purified DNA was quantified using a NanoDrop 2000 (Thermo Fisher Scientific). DNA samples were sequenced using the Myeloid Solution (Sophia Genetics, Boston, MA, USA) and an Illumina Mi-Seq (Illumina, San Diego, CA, USA), using a 2x300 cycle kit. Data files were analyzed using Sophia DDM and Alissa clinical informatics (Agilent, Santa Clara, CA, USA) platforms. Mutations identified in each MV4-11 cell line are listed in Supplementary Table S2.

*Cell growth assay*

Cells were seeded at $1e^5$ cells per mL. Cell number and viability were determined at 0, 24, and 48 h timepoints by trypan blue exclusion (n=3 independent biological replicates) and doubling time calculated as previously described (1). Statistical significance was determined via an unpaired Student's *t*-test with Welch's correction.

*Cell proliferation and apoptosis*

Cell lines were treated with the FLT3 inhibitors sorafenib (Selleckchem), quizartinib (Selleckchem), midostaurin (Selleckchem), and crenolanib (Selleckchem), DNA-PK inhibitor peposertib (Selleckchem), ATM inhibitors KU-60019 (Selleckchem) and WSD-0628 (WayShine Biopharm, Shanghai, China), and N-acetyl-l-cysteine (NAC) (Sigma-Aldrich, Burlington, MA, USA) either alone or in combination. Cells were seeded into 96 well plates at $4e^4$ cells per well, and cell proliferation was measured after 48 h as previously described (1). Half-maximal inhibitory concentration ($IC_{50}$) was determined via non-linear regression analysis and synergy of dose-response and combined effect of the two drugs were assessed using the method of Bliss independence model (4). Cell death was measured using the annexin V/PI fluorescein isothiocyanate/propidium iodide apoptosis and cell viability detection kit (BD Biosciences, SanJose, CA, USA) after 48 h treatment as previously described (5). Data were analyzed and FACs plots generated using FlowJo software version 10.

*Antibodies and immunoblotting*

Protein was extracted from cell lines using RIPA lysis buffer as previously described (5). Proteins were separated on NuPAGE Bis-Tris 4% to 12% gels (Invitrogen, Carlsbad, CA, USA) and transferred onto 0.2 μm nitrocellulose membranes (Bio-Rad, Hercules, CA, USA) for antibody staining (6). Immunoblotting was performed using the following antibodies from Cell Signaling Technology (Danvers, MA, USA) (unless otherwise stated); phospho$^{S2612}$-DNA-PK, total-DNA-PK, phospho$^{S1981}$-ATM, total-ATM, phospho$^{S139}$-H2AX, phospho$^{S392}$-p53, phospho$^{S824}$-KAP1, total-KAP1, total-H3, total-p53 (Abcam, Cambridge, United Kingdom), total-NOX2 (Invitrogen), total-NOX4 (GeneTex, Irvine, CA, USA) and β-actin (Sigma-Aldrich). Secondary antibodies were conjugated with horseradish peroxidase (Sigma-Aldrich). Bands were visualized using a ChemiDoc imager system (Bio-Rad) and data analyzed using ImageLab software.

*Structural modelling of tyrosine kinase inhibitors binding to FLT3*

Several crystal structures detailing the intracellular domains of FLT3, including the inactive conformation of the TKDs, were used as templates to create a multiple sequence alignment homology model of the inactive kinase. The model was generated using Discovery Studio 2018 (Biovia, San Diego, CA, USA), utilizing up to 19 high resolution PDB templates. To construct the kinase in the active conformation, the homologous kinase, DYRK1A in complex with midostaurin (PDB: 4NCT), was used as the sole template. The activation loop (aa826 - aa851) of this active conformation was then grafted onto the ‘inactive’ model, replacing the respective ‘inactive’ activation loop to create a kinase structure representing its catalytically active state. Within this model, the activation loop was then further optimized (in

Discovery Studio, 'loop refinement' and 'minimization' operations) to yield the final 'active' FLT3-TKD model which also represents the active confirmation enforced by the ITD mutation. The binding configurations of sorafenib and quizartinib on each model were investigated using GOLD (Cambridge Crystallographic Data Centre, Cambridge, England). The ChemPLP algorithm was used to rank the stability/affinity of 10 calculated binding poses, with the top poses used to define the protein-ligand interactions.

*pHASED phosphoproteomics*

MV4-11 parental and TKI resistant cells were lysed, reduced, alkylated, and digested in biological triplicates using an optimized protocol as previously described (1). Thereafter, 200 μg of tryptic peptide per sample was used for titanium dioxide phosphopeptide enrichment. Enriched phosphopeptides were lyophilized completely prior to resuspension in 2% ACN/0.1% TFA. Reverse phase nanoflow LC-MS/MS was performed using a Dionex Ultimate 3000RSLC nanoflow high-performance liquid chromatography system coupled with an Orbitrap Exploris 480 MS equipped with a front-end High Field Asymmetric Waveform Ion Mobility Spectrometry (FAIMS) Interface (Thermo Fisher Scientific) as previously described (1).

*Data processing and bioinformatic analysis*

Data analysis was performed using Proteome Discoverer 2.5 (Thermo Fisher Scientific) with previously described dataset parameters (1). Sequest HT was used to search against UniProt *Homo sapiens* database (22,917 sequences, downloaded 22/04/22). Additionally, *Homo sapiens* FLT3 FASTA file containing wild-type and mutant FLT3 sequences (3 sequences, downloaded 21/02/20) were used in this analysis. Fold changes between groups were calculated by Proteome Discoverer using a non-nested pairwise ratio approach, whereby calculated peptide group ratios correspond to the geometric median of all combinations of ratios from all of the replicates for each group (7). Fold changes converted to $log_2$ scale ($log_2$ ratio) by Proteome Discoverer were used for subsequent comparison. Scatter plots were generated using normalized abundances (Supplementary Figure S2), and Pearson correlation was calculated using Perseus (version 1.6.2.2).

*Statistical methods*

Phosphoproteomic data analysis was performed using sensitive and drug-resistant MV4-11 cell lines (n=3 independent biological replicates each). Differentially expressed phosphopeptides and phosphorylation sites were defined as those with a $\log_2$ fold change ≥ 0.5 or ≤ −0.5. Differences between sample groups were analyzed by unpaired Student's *t*-tests and considered significant when $p \leq 0.05$. Graphical data was analyzed and prepared using Perseus (1.6.2.2) (8) and GraphPad Prism (10.0.2) (LaJolla, CA, USA). Results are presented as mean values ± SEM. For *in vitro* (n=3 independent biological replicates) and *in vivo* experiments, graphs were produced using GraphPad Prism Software (version 10.0.2, GraphPad, Boston, MA, USA). Two-sample unpaired Student's *t*-tests with Welch's correction, one-way analysis of variance (ANOVA), or two-way ANOVA were used to determine significant differences between groups unless otherwise indicated. Survival analysis was performed using a log-rank test.

*Ingenuity Pathway Analysis and Kinase-Substrate Enrichment Analysis*

Ingenuity Pathway Analysis software (Qiagen, Hilden, Germany) and Kinase-Substrate Enrichment Analysis (KSEA App, version 1.0) were used as previously described (1).

*Detection of reactive oxygen species*

Dihydroethidium (DHE) (Life Technologies, Australia) was used to detect intracellular cytoplasmic superoxide. Positive controls were treated with 1 mM hydrogen peroxide ($H_2O_2$) for 10 min. Briefly, $5e^5$ cells/mL were washed once in PBS and stained with DHE (adapted from (9)). After staining for 30 min, stain was removed, and cells were resuspended in PBS prior to analysis using a FACSCanto flow cytometer (BD Biosciences). Data was analyzed using FlowJo software (version 10.6.1, BD Biosciences).

*Cell cycle analysis*

MV4-11 FLT3-ITD sensitive and TKI resistant cell lines were maintained as described above. Prior to the start of the experiment, cells were passaged into three independent biological replicates. For FACS analysis, $1e^6$ cells per mL were harvested, washed with PBS, and fixed in ice-cold 75% ethanol prior to incubation at 4°C for 1 h. Cell were then washed to remove ethanol fixation and stained with propidium iodide (PI) (1 µg/mL), followed by the addition of RNase A (0.1 mg/mL). Cells were then incubated in the dark for 30 min prior to analysis using a FACSCanto II flow cytometer (BD Biosciences) The proportional distribution of cells between G0/G1, S, and G2/M phase of the cell cycle was quantified using FlowJo software (version 10.6.1, BD Biosciences), applying the Dean-Jett-Fox model (10).

*Analysis of publicly available patient survival and expression data*

RNA-seq data from the Therapeutically Applicable Research to Generate Effective Treatments (TARGET) and Beat AML initiatives (11,12) was downloaded and TPM values as well as clinical information obtained. Survival analysis was performed using Graphpad Prism. Patients were separated into high and low expression as determined by TPM value for each gene of interest where high expression referred to the top 25% of patients and low expression the bottom 25% of patients (Q1/Q4 split). Cox's proportional hazards regression models were fitted to the survival data. For expression comparison a log2 TPM comparison between groups separated by diagnosis or relapse subtype was conducted in Graphpad Prism. Student's t-test was performed for statistical comparison.

*ATM knockdown*

MV4-11 sensitive and resistant cell lines were transfected using RNAiMAX lipofectamine reagent (Invitrogen, Carlsbad, CA) as per manufacturer's instructions with ATM–specific small interfering RNA (siRNA) at a concentration of 30 nM. The scrambled siRNA control was used at 100 nM. ATM-targeting siRNA was purchased from Thermo Fisher

Scientific (Silencer Select siRNA ID: 111194), and scrambled siRNA control was from Cell Signaling Technologies (#6568) (Danvers, MA).

*Terminal Deoxynucleotidyl Transferase dUTP Nick End Labeling (TUNEL) DNA Fragmentation Assay*

A total of $5e^5$ MV4-11 FLT3-ITD sensitive and TKI resistant cell lines were treated with either 250 nM WSD-0628 for 24 h or 1 mM $H_2O_2$ for 30 min, washed in PBS and fixed in 10% formalin solution (Sigma-Aldrich) for 30 min. Cells were stored in PBS at 4°C prior to staining. For analysis, cells were adhered to pre-coated poly-L-lysine coverslips by placing 50 µL of cell suspension onto the slide and incubating for 30 min at room temperature. Permeabilization of cells was achieved by incubation in PBS containing 0.2% (v/v) Triton-X 100 for 10 min at room temperature before rinsing in PBS. An ApopTag Fluorescein in Situ Apoptosis Detection Kit (cat #S7110; Merck Group, Darmstadt, Germany) was employed to detect apoptotic cells as described by the manufacturer. A negative control sample was prepared by substituting the TdT enzyme for PBS. For a positive control, an additional sample was treated with DNAse 1 buffer (cat #M6101; Promega Madison, WI, USA) for 10 min, followed by another 10 min incubation in DNAse 1 enzyme/buffer at a ratio of 1:1. After ApopTag application, coverslips were washed by submerging three times in PBS, then counterstained with 4′,6-diamidino-2-phenylindole (DAPI; 0.5 µg/mL) for 1 min at room temperature. Coverslips were rinsed three times in PBS before mounting on a slide with Mowiol containing 1,4-diazabicyclo[2.2.2]octane (DABCO). An AXIOplan Imager 2 fluorescence microscope (Carl Zeiss Micro Imaging GmbH, Jena, Thuringia, Germany) was used to count and image all slides. Three independent biological replicates were analyzed per treatment group, with 100 cells counted per sample and the number of TUNEL-positive cells recorded.

*Oxidative DNA damage assessment by 8-hydroxy-2'-deoxyguanosine (8OHdG)*

A total of $5e^5$ MV4-11 FLT3-ITD sensitive and TKI resistant cells were treated with either 250 nM WSD-0628, siRNA targeting ATM or siRNA scramble control (described below). Fixed cells were adhered onto poly-L-lysine coverslips and permeabilized as described above, before blocking by incubation in 50 µL of 1.5% goat serum (v/v) diluted in PBS for 1 h at room temperature. Following three rinses in PBS, DNA/RNA damage antibody (cat #NB110-96878; Novus Biologicals, Centennial, CO, USA) diluted 1:50 in PBS was applied to each coverslip and probed overnight at 4°C. The next day, coverslips were again rinsed three times in PBS before adding the goat-anti mouse AlexaFluor 488 conjugated secondary antibody (cat # A11001; Thermo Fisher Scientific) diluted 1:400 in PBS for 1 h at room temperature in a dark humidified chamber. Coverslips were washed three times in PBS before counterstaining with DAPI for 5 min at room temperature. Following a final wash, coverslips were mounted on slides with Mowiol and viewed with a fluorescence microscope whereby 100 cells were counted per sample and scored positive based on the presence of nuclear fluorescence.

*Type II TKI-resistant luciferase in vivo studies*

All *in vivo* experimental procedures were conducted with approval from the University of Newcastle Animal Care and Ethics Committee (A-2023-308) and performed as previously described (13). Four- to six-week-old immune-compromised NOD-Rag1null IL2rgnull (NRG) mice were purchased from OzGene (WA, Australia) one week prior to xenografting. NRG mice were inoculated with FLT3-ITD sensitive and/or TKI resistant, luciferase positive MV4-11 cells ($1e^6$ cells suspended in PBS) by injection into the lateral tail vein. Bioluminescence imaging (BLI) was used to detect leukemic engraftment, and treatment commenced once BLI reached a mean radiance of $1 \times 10^6$ p/s. Mice were randomized, then treated with either vehicle, sorafenib (10 mg/kg MV4-11 resistant; 2.5 mg/kg MV4-11 sensitive), quizartinib (2 mg/kg) or WSD-0628 (5 mg/kg) as single agents, or WSD-0628 combined with either sorafenib [WSD-0628 (5 mg/kg) + sorafenib (10 mg/kg)], or quizartinib [WSD-0628 (5 mg/kg) + quizartinib (2 mg/kg)] (n=8 mice per group). Treatments were administered once daily. Animal weights were determined twice per week, and systemic leukemic burden was assessed weekly by BLI using an IVIS Spectrum imager (Perkin Elmer), after intraperitoneal injection of luciferin substrate (3 mg per mouse; P1043 Promega). Mice were considered at endpoint and culled if there was weight loss exceeding 20% of initial body weights or body condition scores indicating ethical endpoint. At the end of the 4-week treatment period, 3 mice per group were culled and cardiac blood, spleens, sternum, and femurs were collected for further processing and analysis. For pharmacodynamic assessment, vehicle mice reaching endpoint were given a single dose of treatment (n=2 mice per treatment group) and culled 6 h later for tissue collection.

#### Supplementary Data

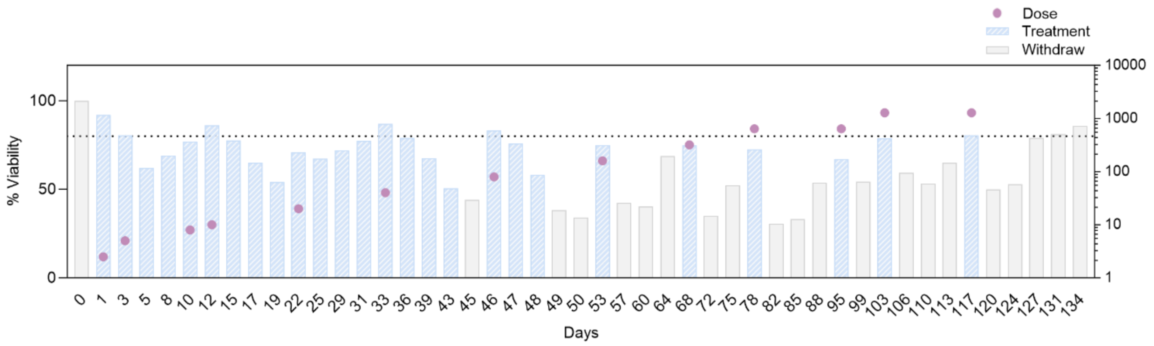


**Supplementary Figure 1. Development of an *in vitro* model of resistance-causing FLT3 mutations**. Resistance was induced by serial passage of MV4-11 parental cell lines in increasing doses of sorafenib over a period of 20 weeks. Sorafenib dose range from 0nM – 1280 nM. Cell viability was assessed via Trypan Blue exclusion. Dotted line= 80% viability

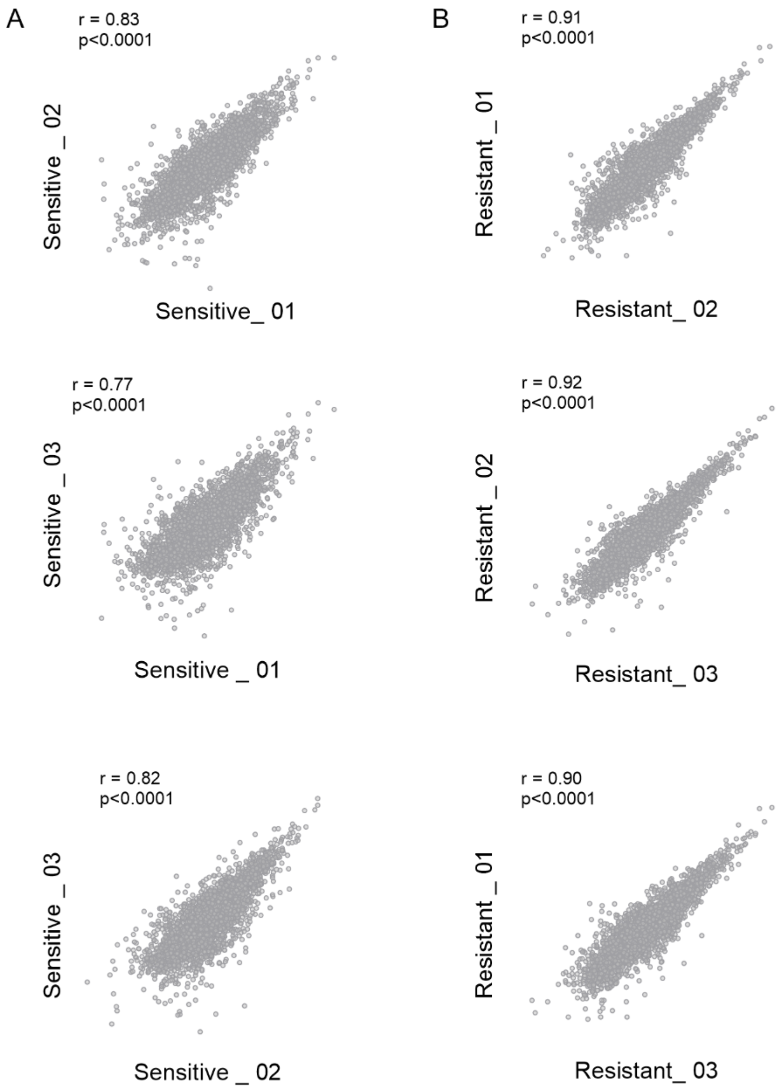


**Supplementary Figure 2. Quantification reproducibility between biological replicates.** Pearson correlation profiles for biological replicates (n=3) of (A) MV4-11 FLT3-ITD sensitive, and (B) MV4-11 FLT3-ITD resistant. Correlation was performed using normalized abundances in Perseus, and graphs were plotted using GraphPad Prism 10.0.2.

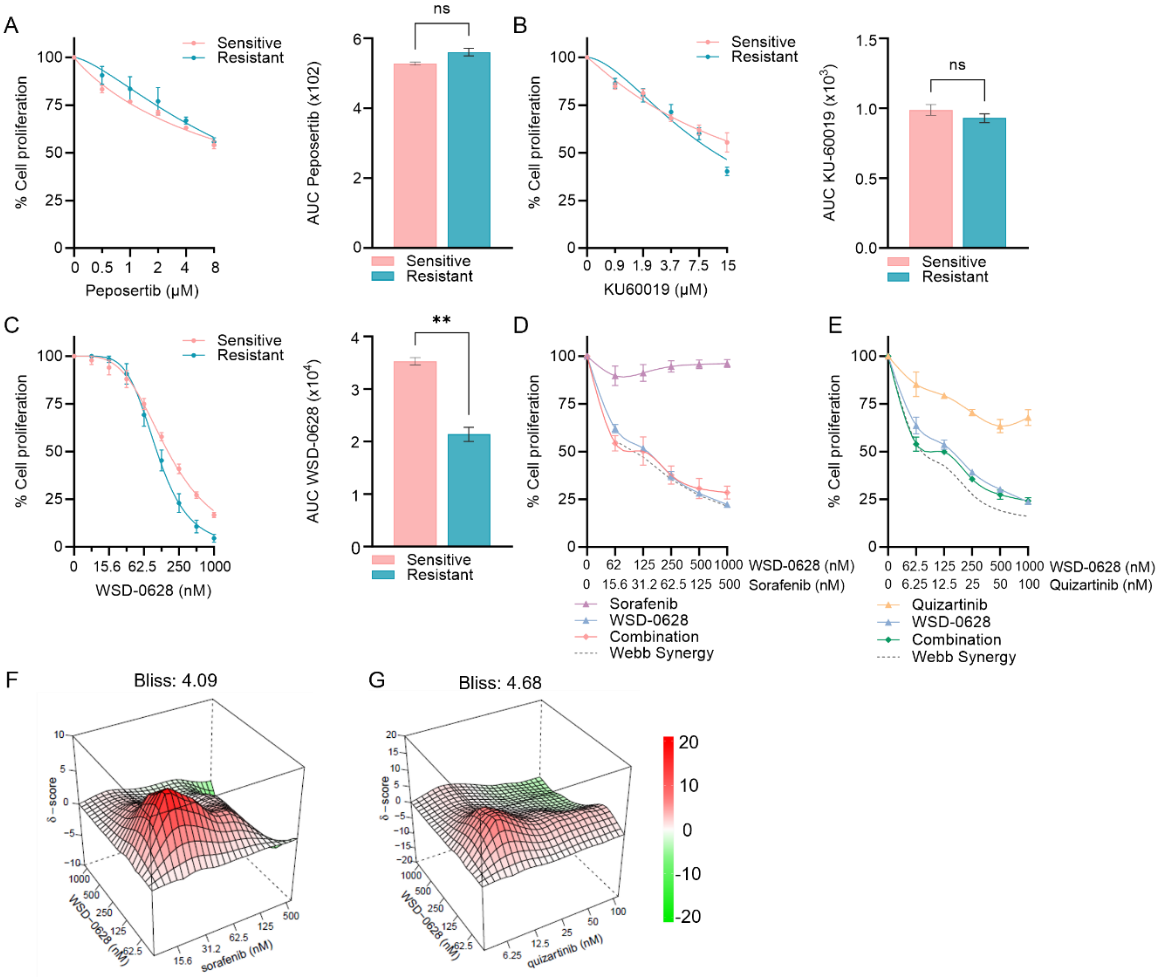


**Supplementary Figure 3. Sensitivity of FLT3-ITD sensitive and FLT3-ITD resistant cell lines to DNA repair inhibitors.** Drug-response was assessed based on cell proliferation assay and area under the curve (AUC) comparison following treatment with the (A) DNA-PK inhibitor peposertib, and ATM inhibitors (B) KU-60019, and (C) WSD-0628 as single agents. Assessment of WSD-0628 in combination with FLT3 inhibitors (D) sorafenib and (E) quizartinib was also performed. Cell proliferation was assessed by resazurin assay at 48-hour treatment. Values shown as mean +/- SEM (n=3 independent replicates). Statistical significance calculated via unpaired t-test with Welch's correction. Significance threshold of *$p$<0.05, **$p$<0.01 and ***$p$<0.001. Bliss synergy analysis of combined effect of WSD-0628 with (F)

sorafenib or (G) quizartinib in FLT3-ITD resistant cell lines (<0= antagonistic, >0<10=additive, >10=synergistic).

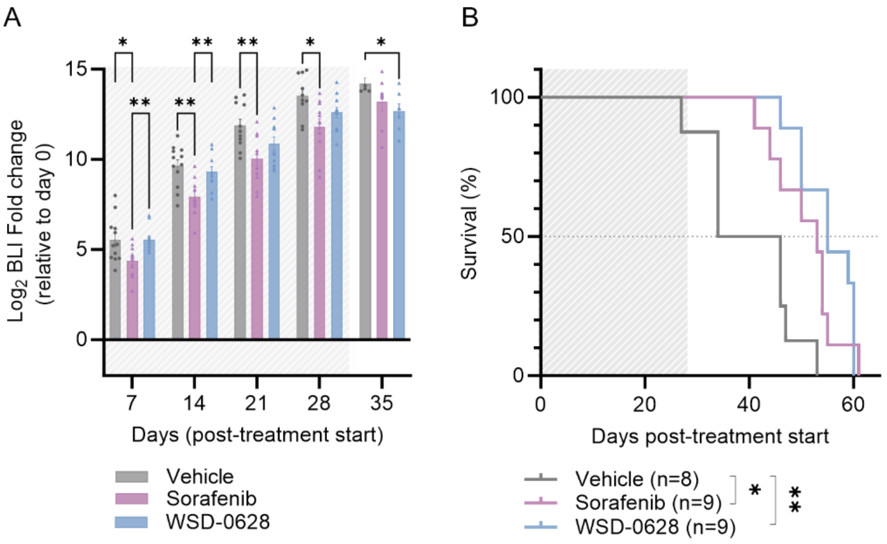


**Supplementary Figure 4. In vivo sensitivity to ATM inhibition in MV4-11 FLT3-ITD cell lines sensitive to tyrosine kinase inhibitor (TKI) therapy.** MV4-11 FLT3-ITD cell lines sensitive to TKI were injected into the lateral tail vein of NOD-Rag1null IL2rgnull (NRG) mice. Treatment commenced once BLI reached a mean radiance of 1 × $10^6$ p/s. Sorafenib, and WSD-0628 were administered once daily for 4 weeks. (A) Monitoring of leukemia burden using bioluminescence BLI imaging over time (shaded area indicates treatment time) of mice treated with WSD-0628, or sorafenib. (B) Kaplan-Meier survival analysis of MV4-11–Luc+ FLT3-ITD sensitive cells (n= 8-9 mice per group, shaded area indicates treatment time) treated with WSD-0628, or sorafenib. Log-rank (Mantel-Cox) Test; Significance threshold of *$p$<0.05, **$p$<0.01 and ***$p$<0.001.
